# An aqueous watercress extract with topical anti-inflammatory properties in inflamed human skin

**DOI:** 10.1101/2025.09.09.675161

**Authors:** Paul G Winyard, Darren Phillips, Hannah Folland, David Caballero-Lima, Bartosz Cwikla, A Toby A Jenkins, Kyle Stewart

## Abstract

Watercress (*Nasturtium officinale*) is a species of the Brassicaceae plant family, with a biochemical composition which suggests potential uses in anti-inflammatory skincare applications. Watercress has a high content of both anti-inflammatory compounds (*e.g*., polyphenols) and compounds which upregulate the Nrf2-mediated antioxidant gene response (*e.g.*, isothiocyanates) in human tissues. We have developed a process to manufacture an aqueous watercress extract that contains isothiocyanates and flavonoid-type polyphenols, some of which are known anti-inflammatory compounds. We partially characterized the chemical composition of our extract (LC-QTOF-MS), and we demonstrated in a cultured full-thickness human skin model, that the extract was non-cytotoxic and non-irritant, as monitored by cell viability and IL1α concentrations, respectively. In human trials, *in vivo*, we showed by visual clinical scoring of inflammation that the extract had anti-inflammatory properties. The anti-inflammatory properties of the aqueous watercress extract may provide a treatment for inflammatory skin diseases such as eczema.

## 1. Introduction

Watercress, *Nasturtium officinale,* is an aquatic perennial plant that belongs to Brassicaceae family. It is a leafy green vegetable commonly consumed by humans, and medicinal properties have been attributed to this plant (Klimek-Szczykutowicz et al, 2018; Kokhdan et al, 2021). Watercress contains bioactive phytochemicals with potential anti-inflammatory (Shahani et al, 2017) and antioxidant (Zeb, 2015) properties. There is interest in using natural plant products in topical anti-inflammatory applications for the skin, due to relatively low cytotoxicity (Fernandes et al, 2023). The antioxidants in watercress include carotenoids, chlorophyll and other polyphenolic compounds (Zeb, 2015). Watercress also contains water-soluble glucosinolates (S-glucopyranosyl thiohydroximates) which are enzymatically hydrolysed to isothiocyanates (e.g. phenethyl isothiocyanate; PEITC) when the plant tissue is macerated (Rose et al, 2000; Kyriakou et al, 2022; Kyriakou et al 2023; Kyriakou et al, 2024). The mechanism of isothiocyanate production involves the disruption of the plant tissue by maceration, causing the mixing of glucosinolates with the plant enzyme, myrosinase. In this way, the enzyme catalyses the hydrolysis of glucosinolates to isothiocyanates. Following watercress ingestion, isothiocyanates induce phase II enzymes in humans (Rose et al, 2000). Additionally, some of the key individual compounds from watercress which reportedly have anti-inflammatory activities include a range of isothiocyanates (Rose et al, 2005a; Tsai et al, 2010). Isothiocyanates activate the human transcription factor, Nrf-2 (Ernst et al, 2013) which, in turn, upregulates the expression of certain antioxidant enzymes, including catalase and haem oxygenase (HO-1). HO-1 exerts anti-inflammatory effects by removing the pro-oxidant, haem, and by catalysing the generation of the anti-inflammatory products, carbon monoxide and biliverdin (Saha et al, 2020). It appears that HO-1 also exerts anti-inflammatory activities through other mechanisms, such as macrophage polarisation (Ryter et al, 2021).

The above mechanisms appear to be associated with the anti-inflammatory activity of watercress: ingestion of watercress extracts had anti-inflammatory effects in rodents (Sadeghi et al, 2014; Shahani et al, 2017), whilst a systematic review concluded that ingestion of watercress extracts or raw watercress had anti-inflammatory effects in humans (Wen et al, 2025). A study by Schulze et al (2021) showed that human consumption of raw watercress caused an initial pro-inflammatory response followed by an increased exercise-induced response in terms of both “pro-inflammatory” and “anti-inflammatory” cytokines. Camponogara et al (2019) reported that the topical application of an extract of watercress caused a reduction in croton oil-induced skin inflammation in a mouse model. The latter study provided evidence that the anti-inflammatory activity of the extract was mediated through pathways involving the transcription factor, NF-κB, and the glucocorticoid receptor. However, there are limited reports regarding the anti-inflammatory effects of watercress extracts on human skin when applied topically. Furthermore, a need remains for the development of efficient and scalable processes for the recovery of bioactive compounds from plants such as watercress. For some applications, especially those related to human consumption or topical application, a watercress extract which is aqueous, avoiding the use of organic solvents, is advantageous when considering toxicity and cost. The present paper reports: (1) a method suitable for the medium-scale manufacturing of a stable aqueous extract from watercress, (2) chemical characterisation of some of the constituent compounds of the extract, (3) an assessment of the cytotoxicity/irritancy of the extract in an *in vitro* model of full thickness human skin (a living dermal equivalents model known as Labskin^1.1 ©^) and (4) an *in vivo* demonstration of bioactivity of the extract as an anti-inflammatory agent when topically applied to inflamed human skin.

## 2. Methods

### 2.1. Sample preparation

The fresh watercress used in this trial was obtained from The Watercress Company (Dorchester, Dorset, UK). The supplied plant material (10 Kg of dark green variety) was harvested to a specification of 20 cm length. Watercress was refrigerated in a cold room at 4°C for 24 h before the extraction process began.

The watercress was prepared using a RoboQbo 15-4 food processor (Roboqbo Srl, Italy), which has a sample bowl equipped with a cutting blade. Using a cutting blade, speed and residence time were investigated to determine the optimum conditions to homogenise the material. The bowl was loaded with 0.5 or 1.0 Kg of watercress and chopped for 5 min at 2000 rpm. Using these cutting conditions, samples of the homogenised mixture were taken at time intervals of 0 min (immediately after cutting), 5 min, 10 min, 30 min, 1 h, 2 h, 3 h, 6 h and 7 h. Samples were collected and sealed in a 50 mL sample tube then immersed in a water bath at 80°C for 1 h. The standard procedure was that samples were then centrifuged at 14,000 *x **g*** for 10 min and filtered through a 0.25 µm PTFE syringe filter. To test the effects of variations in the last 2-3 steps of the above protocol (*i.e.*, different centrifugation and filtering conditions, along with the effects of autoclaving), four different versions of the extract were prepared, as follows. Extract A: centrifuged (10,000 *x **g*** for 10 min), autoclaved (121 °C for 15 min). Extract B: centrifuged (10,000 *x **g*** for 10 min), filtered (0.25 µm). Extract C: centrifuged (10,000 *x **g*** for 10 min), autoclaved (121 °C for 15 min), centrifuged (10,000 *x **g*** for 10 min). Extract D: centrifuged (10,000 *x **g*** for 10 min), autoclaved (121 °C for 15 min), filtered (0.25 µm). The filtrates were collected and 1 mL aliquots transferred to Eppendorf tubes and stored at -20°C before analysis.

### 2.2. Analysis of watercress extract by UHPLC-QTOF

The analysis of the extracts was performed with an Agilent 1290 Infinity ultra-high performance liquid chromatography (UHPLC) system equipped with an UV detector and coupled with detection by quadrupole time-of-flight (QTOF) mass spectrometry. The charged ions, as generated by electrospray ionisation (ESI), were detected as either positively charged ions (Supplementary Table 1) or negatively charged ions (Supplementary Table 2) by use of the instrument in positive ion mode or negative ion mode.

Chromatographic separation was achieved using a Waters Acquity BEH C18 (1.7 µm x 2.1 mm x 100 mm) column, with a sample injection volume of 1.0 µL. A mobile phase of 1% v/v methanol/water (0.1% v/v formic acid in both solvents) was used at a flow rate of 0.25 mL/min, with the column temperature maintained at 30°C. The UHPLC was coupled to an Agilent 6530 Accurate-Mass Q-TOF LC-MS instrument. Mass spectra were collected following ESI in both negative and positive ion mode of the eluent. Spray chamber conditions were: gas temperature 250°C, drying gas flow, 8.0 L/min, nebuliser pressure, 30 psig; capillary voltage (Vcap), 4500V; nozzle voltage, 500 V; fragmentor voltage, 150 V; skimmer voltage, 65 V. Elution was monitored by UV absorbance at 254 nm, and mass spectra recorded in the mass range m/z 60-1500. Data analysis was performed using Agilent Masshunter Software. Components were identified using extracted ion chromatogram (EIC) scans.

### 2.3. Cytotoxicity and skin irritation assessment of the watercress extract

*N. officinale* aqueous extracts were prepared, including four variations in the final 2-3 steps of the protocol (as described in section 2.1) to produce four versions of the extract: A, B, C and D. These four extracts underwent subsequent *in vitro* testing of skin cytotoxicity. This study was conducted at the facilities of Labskin Ltd. An *in vitro* model of human skin was prepared (Lewis et al, 2018); primary adult human dermal fibroblasts were embedded into a fibrin matrix to produce dermal equivalents (DEs). DEs were cultured to allow the fibroblasts to remodel the matrix. Primary neonatal human keratinocytes were applied to the dermal equivalent surface and cultured under liquid for 48 h. The cells were cultured at the air-liquid interface until a stratified epidermis was formed. Incubation conditions for all cultures were 37 ± 2 °C in 5 ± 1 % (v/v) CO_2_ at ≥ 95% relative humidity. The resulting skin constructs were used as an *in vitro* model of human skin, known as Labskin^1.1©^. The cytotoxic and irritant effects of the aqueous watercress extracts were tested in comparison with the known skin irritant, sodium dodecyl sulfate (SDS). Human skin equivalents (HSEs; in quintuplicates for each treatment) were treated with 11 µL of each test item (Extracts A, B, C or D) or control. Dulbecco’s phosphate-buffered saline (dPBS) was the non-irritant negative control and SDS was an irritant positive control. All HSEs were exposed to the test items/controls for 20 ± 1 min at 37 ± 2 °C in 5 ± 1 % (v/v) CO_2_ at ≥ 95% relative humidity. After 20 min, test items were rinsed off each HSE with dPBS and the stratum corneum surface was dried with sterile filter paper. HSEs were then incubated at 37 ± 2 °C in 5 ± 1 % (v/v) CO_2_ at ≥ 95% relative humidity for an additional 24 ± 2 h.

Following the 24 ± 2 h post-treatment incubation, assessments were performed using the following end points: (1) The undernatant was aspirated and frozen for quantification of cytokines by ELISA using the R&D systems Quantikine® ELISA for human interleukin-1α (IL-1α/IL1F1), according to the manufacturer’s instructions. (2) The cytotoxicity of irritant compounds was assessed by quantification of cell viability in dermal equivalents using an MTT (3-(4,5-dimethylthiazol-2-yl)-2,5-diphenyltetrazolium bromide) assay. To do this, the undernatant was replaced with a 1 mg/mL solution of MTT in Labskin maintenance medium. The HSEs were incubated at 37 ± 2 °C in 5 ± 1 % (v/v) CO_2_ at ≥ 95% RH for 3.5 h. The formazan produced was extracted using 2-propanol puriss (ACS reagent, >99.8%, Sigma-Aldrich) by immersing Labskin constructs in it overnight. Extracted formazan was quantified using a spectrophotometer at a wavelength of 540 nm. The absorbance readings obtained were used to calculate the mean average and normalise to the negative control (dPBS) to provide the results expressed as percentage cell viability. The above protocol follows the OECD guidelines for testing chemicals for skin irritation (OECD, 2025).

### 2.4. Testing of the anti-inflammatory activity of the watercress extract in an *in vivo* human model of skin inflammation

Aqueous watercress extract samples were produced in accordance with the standard sample preparation method described above, using a sample of the homogenised mixture taken at 2 h after cutting. The samples were entered into a single-blind, within-subject clinical trial, with accreditation according to ISO 9001:2015 and ISO 17025:2017. The study was overseen by a consultant dermatologist. A 96-h human patch test in 30 male and female subjects was undertaken to investigate the effect of the extract on skin irritation when administered with a known skin irritant (positive control – SDS, 0.3% w/v). All subjects were healthy, consenting adults over the age of 18 of either sex. There were no exclusions or withdrawals of test subjects. The age range was 19-71 with 20 female and 10 male subjects and a Fitzpatrick Skin Assessment Range of 9-24. The study was performed in accordance with the principles of Good Clinical Research Practice and in consideration of the Helsinki Declaration (Helsinki Declaration 64th WMA General Assembly, Fortaleza, Brazil, October 2013) relating to ethical principles for research involving human subjects. It followed the COLIPA 1997 “Guidelines for the Assessment of Skin Tolerance of Potentially irritant Cosmetic ingredients” (Walker et al, 1997). Subject data were held according to the requirement of the UK General Data Protection Regulation (GDPR) with the data being archived electronically. The data were anonymised.

The study applied the following Exclusion Criteria: pregnant women; subjects with excessive blemishes, marks, scars, sunburns on the test site(s) which could interfere with scoring; forms of medication which may affect skin response; signs of pre-existing skin irritation on test site(s); participation in simultaneous studies which that might interfere with the test evaluation; minors. During the study the following withdrawal criteria were applied: subjects who did not follow the requested conditions; subjects who suffered any illness or accident or developed any condition which could affect the outcome of the study; subjects who no longer wished to participate in the study.

Subjects were asked to refrain from exposure to UV rays and to avoid getting the patches wet and asked to provide information concerning the use of any drug, with particular reference to anti-inflammatory drugs, steroids and antihistamines. Test items (0.06 mL) were applied under occlusive dressings over a 96-h period with a wipe-off and re-apply regime every 24 h (four applications). The patches consisted of an occlusive Finn Chamber on Scanpor 8 mm tape to which Webril (Kendall Corporation) discs, approximately 2.5 cm in diameter, were fixed along the midline and applied mainly on the upper back, or upper arms when the upper back was not suitable. The area of skin designated for patch testing was cleaned with demineralised water and towel dried. The patch was applied to the cleaned area.

At the applicable time-intervals the patch was removed, and residue was wiped away. Approximately 15 min after removal of the patch the application area was carefully examined by the competent study monitor to evaluate skin reactions and grade accordingly with values ranging from 0 to 3 (see below) to express differences in observable reactions. The negative control was deionised water, and the positive control was SDS (0.3%). The sites were assessed by the same qualified assessor on all days according to the European Society of Contact Dermatitis guidelines for diagnostic patch testing (Johansen et al, 2015). Illumination of the sites for assessment was by a 60 W pearl bulb, approximately 30 cm away from the site. Skin reactions were scored at the first, second, third and fourth dressing removals, according to the Evaluation Scale (which includes erythema, oedema, dryness/desquamation and vesicle size/visibility. Mean Daily Irritation Scores (MDIS; with permitted integer values of 0 to 3 inclusive, where 0 = “none” and 3 = “severe”) were calculated by adding up the evaluation grades recorded for the subjects within the panel, then dividing the total by the number of subjects. A mean value was therefore used.

### 2.5. Statistical analysis

Data handling, statistical analysis and data representation was carried out using GraphPad Prism v. 9.4.1. Methods of calculations are indicated in the text and a p-value of ≤ 0.05 was considered statistically significant.

## 3. Results

### 3.1. Analysis of aqueous watercress extract by UHPLC-QTOF-MS

The aqueous extraction process resulted in light brown solutions with pH values of 5.5-6.2. The described method resulted in the visible removal of both the insoluble plant fibre and the heat-precipitated protein fraction, during the centrifugation step of the process. A range of constituent chemical compounds of interest were identified in the aqueous extract by UHPLC-QTOF mass spectrometry. A compilation of the mass spectrometry analysis of the extract’s composition - generated in positive and negative mode - is shown in Table 1. These analytes were screened in positive and negative modes and tentatively identified using the EIC function. Positive and negative mode results are separately shown in Supplementary Tables 1 and 2. For examples of the observed EICs, see Figure 1.

**Table 1.** Analysis of watercress aqueous extract: compilation of UHPLC-QTOF mass spectrometry data generated in positive and negative modes. (+) Tentative assignment, based on chromatographic elution time and mass spectral analysis, (↑) Concentration increasing with time, (↓) Concentration decreasing with time.

| Component | Molecular formula | Ionisation | Retention time (min) | Timepoint |  |  |  |  |  |  |  |  |
| --- | --- | --- | --- | --- | --- | --- | --- | --- | --- | --- | --- | --- |
|  |  |  |  | 0 | 5 min | 10 min | 30 min | 1 h | 2 h | 3 h | 6 h | 7 h |
| Glucobrassicin and derivatives |  |  |  |  |  |  |  |  |  |  |  |  |
| 1-Methoxyglucobrassicin | C17H22N2O10S2 | Negative | 4.7 | (+) | (+) | (+) | (+) | (+) | (+) | (+) | (+) | (+) |
| 4-Methoxyglucobrassicin | C17H22N2O10S2 | Negative | 4.7 | (+) | (+) | (+) | (+) | (+) | (+) | (+) | (+) | (+) |
| 4-Hydroxyglucobrassicin | C16H20N2O10S2 | Both | 5.2 | (+) | (+)(↑) | (+)(↑) | (+)(↑) | (+)(↑) | (+)(↑) | (+)(↑) | (+)(↑) | (+)(↑) |
| Isothiocyanates and isothiocyanate reaction products |  |  |  |  |  |  |  |  |  |  |  |  |
| S-(N-β-phenylethylthiocarbamoyl)glutathione (PTCG) | C19H26N4O6S2 | Positive | 6.0 | (+) | (+) (↓) | (+) (↓) | (+) (↓) | (+) (↓) | (+) (↓) | (+) (↓) | (+) (↓) | (+) (↓) |
| Phenethyl isothiocyanate (PEITC) | C9H9NS | Negative | 3.5 | (+) | (+) (↑) | (+) (↑) | (+) (↑) | (+) (↑) | (+) | (+) | (+) | (+) |
| 7-Methylsulfinylheptyl isothiocyanate | C9H17NOS2 | Positive | 7.9 | (+) | (+) (↑) | (+) (↑) | (+) (↓) | (+) (↓) | (+) | (+) | (+) (↓) | (+) (↓) |
| 8-Methylsulfinyloctyl isothiocyanate | C10H19NOS2 | Positive | 8.6 | (+) | (+) (↑) | (+) (↓) | (+) (↓) | (+) (↓) | (+) (↓) | (+) (↓) | (+) (↓) | (+) (↓) |
| 4-Methylsulfinylbutyl isothiocyanate (sulforaphane) | C6H11NOS2 | Positive | 2.8 | (+) | (+) ↑ | (+) ↑ | (+) ↑ | (+) ↑ | (+) ↑ | (+) ↑ | (+) ↑ | (+) ↑ |
| Other components |  |  |  |  |  |  |  |  |  |  |  |  |
| Tryptophan | C<sub>11</sub>H<sub>12</sub>N<sub>2</sub>O<sub>2</sub> | Both | 3.9 | (+) | (+) | (+) | (+) | (+) | (+) | (+) | (+) | (+) |
| Patchouli alcohol | C<sub>15</sub>H<sub>26</sub>O | Positive | 3.7 | (+) | (+) ↑ | (+) ↑ | (+) ↑ | (+) ↑ | (+) ↑ | (+) (↓) | (+) (↓) | (+) (↓) |

**Figure 1.**
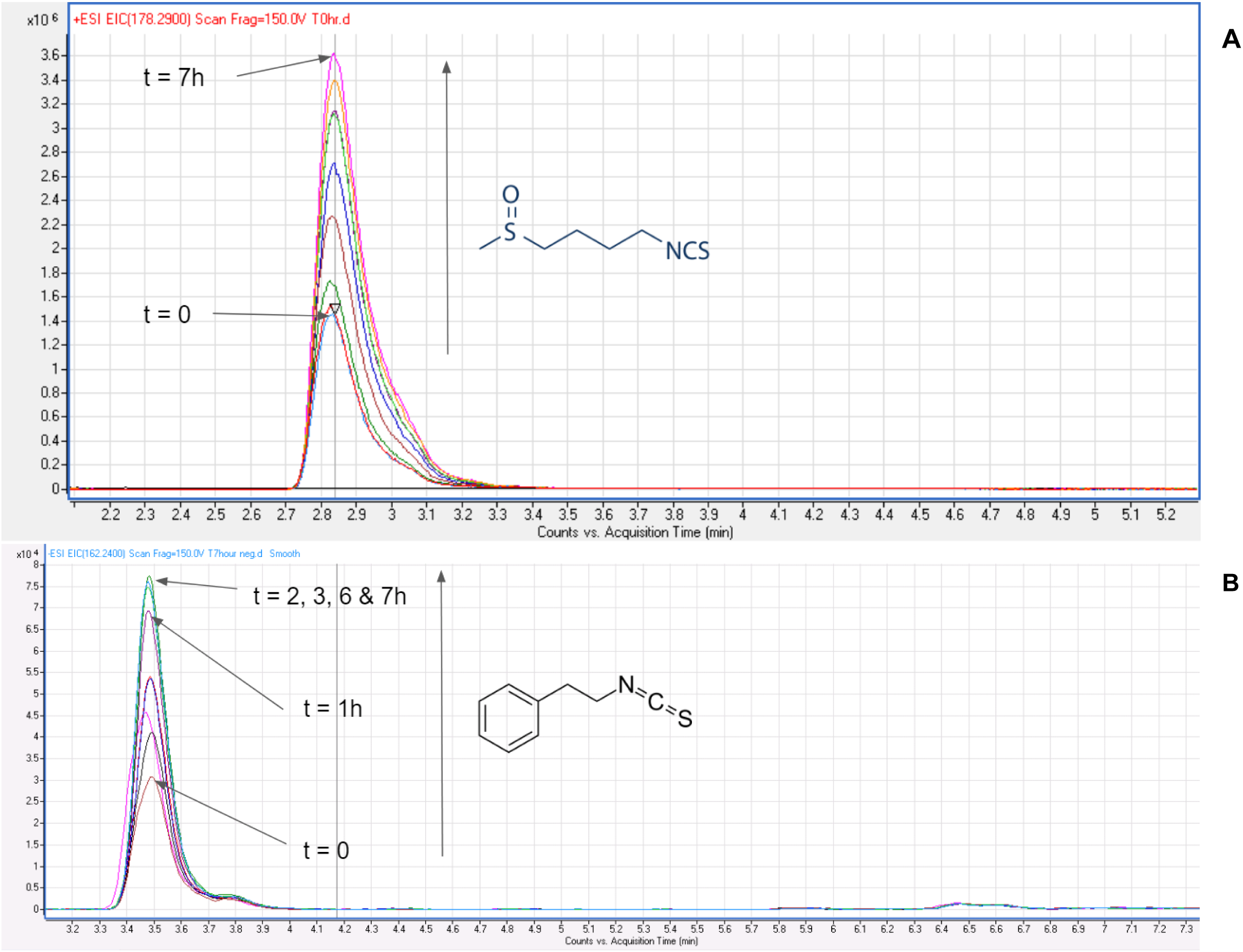
Extracted ion chromatograms (EICs) from UHPLC-QTOF mass spectrometry analyses in the positive mode. Panel **A:** the ion at m/z = 178.29 is putatively assigned to sulforaphane. Panel **B**: the ion at m/z = 162.24 is putatively assigned to PEITC.

The peak heights of some of the detected compounds within the watercress aqueous extract were found to vary with time. Examples of this are shown in Fig. 1, being the cases of the putatively assigned peaks of the isothiocyanates, 4-methylsulfinylbutyl isothiocyanate (sulforaphane) and phenethyl isothiocyanate (PEITC). The change in peak intensity is shown in Table 1, with indicated increases (↑) and decreases (↓) over time, relative to the peak area of the previous time point. It was found that the peak intensity of the sulforaphane product ion, [M+H]^+^, at 178.29, increased over the course of 7 h (Fig. 1), suggesting the concentration of sulforaphane increases over this time. The putatively assigned component of the aqueous watercress extract mentioned above, PEITC, is a widely reported compound in watercress. As shown in Fig. 1, it was found that the extracted ion, [M-H]^-^ at 162.24, also increased over the time-period from t = 0 to t = 2 h, after which the peak intensity plateaued and remained relatively constant.

### 3.2. Cytotoxicity and irritancy assessment of the *N. officinale* extract in an *in vitro* 3D model of full thickness human skin

Labskin^©^ is an *in vitro* 3D model of full thickness human skin (Lewis et al, 2018). *N. officinale* extract was tested, following the OECD guidelines for testing chemicals for skin irritation: cell viability testing, and additionally, the effect on a pro-inflammatory cytokine, IL-1α (OECD, 2021). The final centrifugation/filtration/autoclaving steps in the preparation of the *N. officinale* extracts were varied in four different ways (A, B, C, D) – see section 2.1. Extracts B, C and D did not affect cell viability compared to the negative control (Dulbecco’s phosphate buffer saline (dPBS); Fig. 2, panel A). The application of extract A, in the Labskin model, caused a small but statistically significant (p=0.05) increase in cell viability. The pro-inflammatory cytokine, IL-1α, is produced after a chemical irritant is placed on the skin (Cavalli et al, 2021). Exposure to the chemical irritant, SDS, induced a 48-fold increase in the IL-1α concentration compared to the negative control dPBS (*p*<0.0001, Tukey’s multiple comparisons test). *N. officinale* extract did not significantly affect the IL-1α production when compared to dPBS (Fig. 2, panel B). Results from this *in vitro* model of human skin suggest that the presence of the *N. officinale* extract did not decrease skin cell viability and did not induce an inflammatory response.

**Figure 2.**
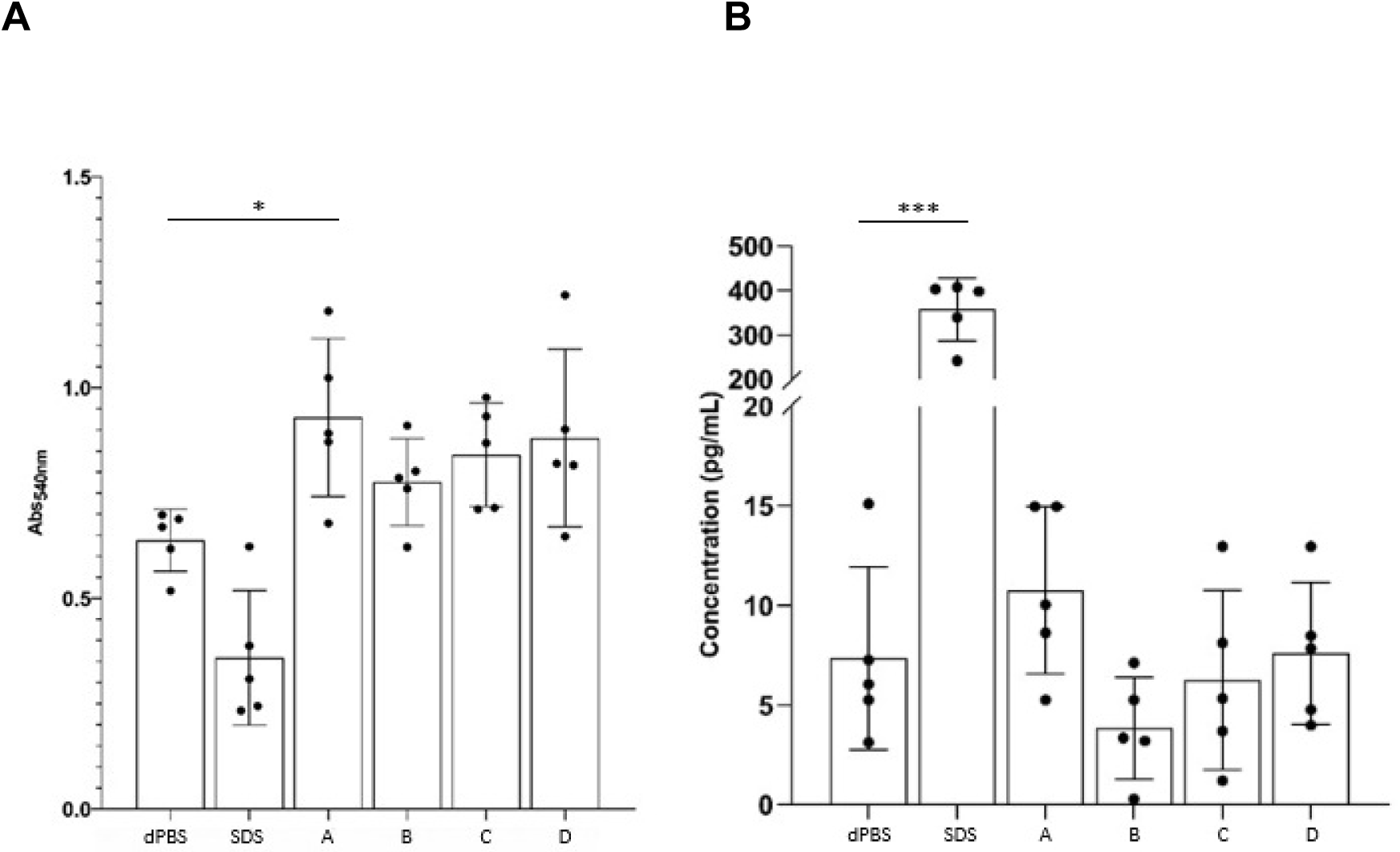
Graphs showing the results from the testing of *Nasturtium officinale* extract in relation to percentage cell viability and irritation (as assessed by interleukin-1α release) using an in vitro full thickness human skin model (Labskin^1.1^ ^©^ dermal equivalents; DEs). Panel **A**: Cell viability in DEs measured using the MTT assay, in which the chemical reduction of MTT was measured as absorbance at 540 nm (A_540_). Four different preparations of the *Nasturtium officinale* extract were compared to a negative control (Dulbecco’s phosphate buffered saline; dPBS) and a positive control (sodium dodecyl sulfate (SDS); 5%, w/v). Panel **B**: Concentration of interleukin-1α produced from DEs, upon exposure to dPBS, 5% SDS, or *N. officinale* extracts. The data were analyzed using an Ordinary One-Way ANOVA: *p* = 0.05 (*); *p* < 0.0001 (***). N = 5 in each group. Error bars indicate ± SD.

### 3.3. Anti-inflammatory activity of the watercress extract in SDS-driven human skin inflammation *in vivo*

Both the neat extract alone, and the 10% extract alone, resulted in a skin irritation score (MDIS) of 0.00 at 24, 48, 72 and 96 h, with no evidence of any inflammatory response in any test subjects across the duration of the trial (data not shown). Fig. 3, panel A, shows that when the extract, diluted to 10%, was co-applied with the 0.3% SDS irritant there was a reduction in irritation score. Fig. 3, panel B, shows the effect when the extract was applied at the 48-hour time-point, highlighted by the arrow, when irritation had already become established. When the extract was applied under these experimental conditions, there was a statistically significant lowering of skin irritation at the 72 and 96 h time-points.

**Figure 3.**
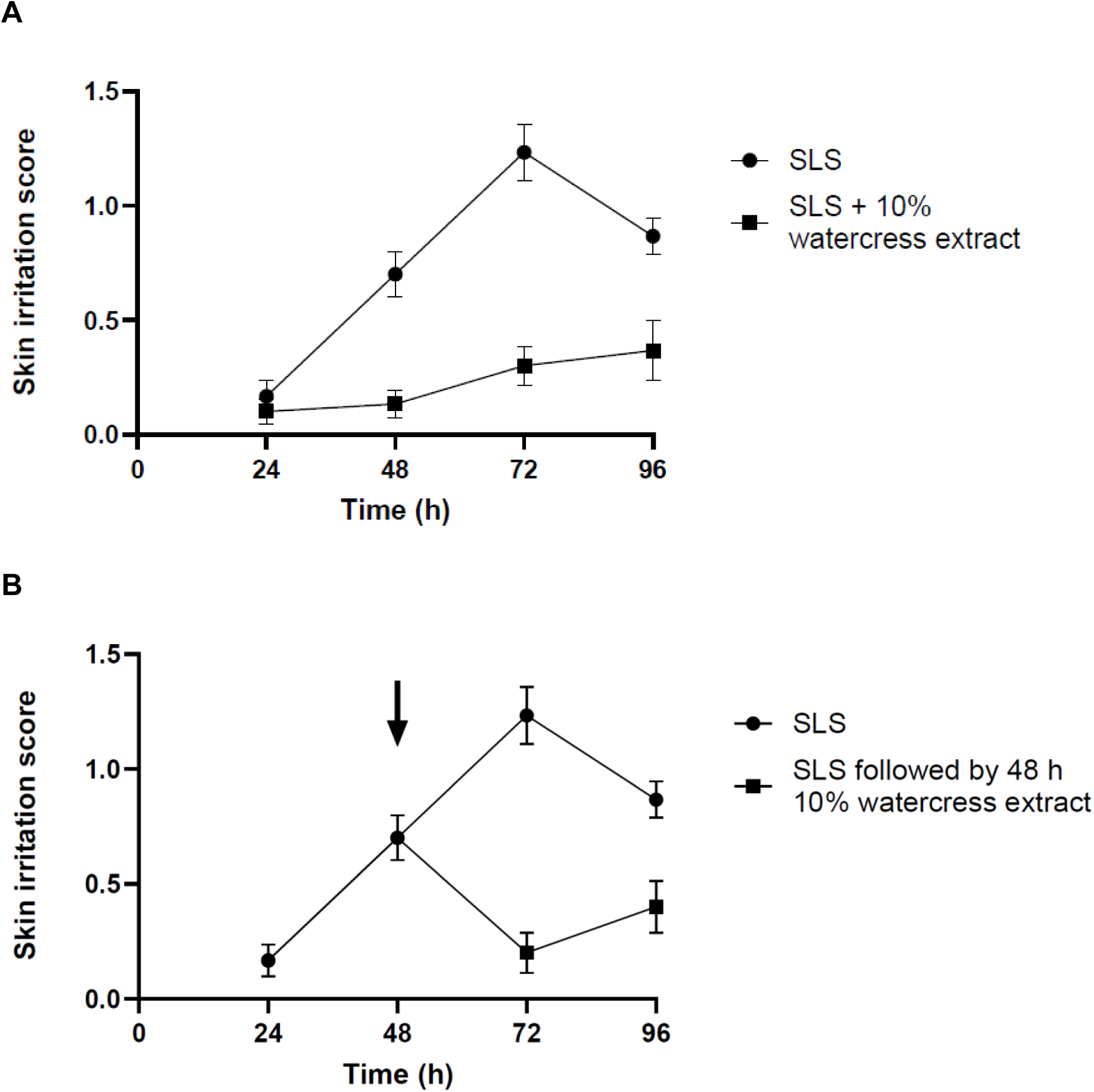
*In vivo* effect of watercress extract on SDS-induced human skin inflammation. Panel **A**: effect of watercress extract (10%) on SDS-induced human skin inflammation, using a protocol where either SDS alone, or SDS mixed with the extract, were repeatedly topically applied (every 24 h) to the skin. Panel **B**: effect of watercress extract (10%) on SDS-induced skin inflammation, using a protocol where SDS alone was topically applied at the t = 0 and t = 24-h time-points, followed by (beginning at the 48-h time-point) the repeated application (every 24-h) of either SDS alone, or SDS mixed with the extract. N = 30 in each group. Error bars indicate ± SEM.

## 4. Discussion

The current study reports a method for obtaining an aqueous extract from the leaves and stems of fresh watercress, avoiding the addition of organic solvents. Such an approach to extracting bioactive compounds from watercress has the advantages of (a) sustainability and (b) avoidance of the potential toxicity of organic solvents. The extraction process includes an incubation period following the cutting of the watercress plant tissue, to allow the enzyme myrosinase (released from the macerated plant tissue) to catalyse the generation of isothiocyanate products from glucosinolates within the mixture. A heating step at the end of the incubation period results in the precipitation of proteins and the inactivation of myrosinase. The precipitated protein fraction, together with the plant fibre, is removed by centrifugation and/or filtration. The aqueous extract of watercress was characterised as containing potentially bioactive compounds, as will be further discussed below.

The resulting aqueous extract was subjected to LC-MS analysis to provide a partial chemical characterisation of the extract’s constituents. We mainly focused on analysis of glucosinolates and their enzymatic hydrolysis products, isothiocyanates, as these compounds are thought to underlie some of the key bioactivities of watercress extracts (Rose et al, 2000, Rose et al 2005a; Rose et al 2005b; Kyriakou et al 2022, Kyriakou et al, 2023; Kyriakou et al, 2025). Several glucosinolates and several isothiocyanates were detected in the present study. The detected glucosinolates included 1-methoxyglucobrassicin, 4-methoxyglucobrassicin and 4-hydroxyglucobrassicin. The detected isothiocyanates – which are products of the hydrolysis of glucosinolates – included PEITC, 4-methylsulfinylbutyl isothiocyanate (sulforaphane), 7-methylsulfinylheptyl isothiocyanate and 8-methylsulfinyloctyl isothiocyanate. The presence of several of these glucosinolates and isothiocyanates in watercress extracts has previously been reported (Rose et al, 2000; Rose et al 2005b; Schuchardt et al, 2019). PEITC readily reacts with nucleophilic groups, and a β-phenylethyl-glutathione conjugate, *S*-(*N*-β-phenylethylthiocarbamoyl)glutathione (PTCG), was also detected. This conjugate was also previously detected in watercress extracts (Rose et al, 2000). Although organic solvents such as methanol and ethanol provide efficient extractions of some phytochemicals from watercress, aqueous extracts also provide significant recoveries of some isothiocyanates and polyphenols from plant tissues (Talalay et al, 2007; Sengupta et al, 2025).

Having ascertained the presence in the extract of potentially bioactive compounds, we aimed to establish whether the extract exhibited toxicity to human skin and whether the extract possessed anti-inflammatory properties. To test the toxicity of the extract, we employed an *in vitro* full thickness skin model (“Labskin” HSEs). HSEs recapitulate key aspects of human skin biology (Lewis et al, 2018). SDS (also known as sodium lauryl sulfate) is a commonly employed chemical irritant for inducing a loss of skin barrier integrity and an inflammatory response (James-Smith et al, 2011). *In vitro*, exposure of dermal equivalents to SDS (at a concentration up to 5%) results in a loss of viability (as monitored by MTT reduction; OECD, 2025) and/or an increase in IL-1α release (Olsen et al, 2018) in reconstructed models of the human epidermis. In contrast, in the present study there were no overt effects of the undiluted watercress extract on MTT reduction or IL-1α levels, suggesting that the extract was non-toxic under these experimental conditions.

Having established the apparent absence of toxicity of the aqueous watercress extract, we next progressed to testing the bioactivity of the extract in terms of its effects on inflammation in a human *in vivo* patch-test-based model of skin inflammation. We investigated the capacity of the extract to prevent inflammation when co-administered with a known skin irritant, and the capacity of the extract in treating established irritation caused by a known skin irritant. Again, the irritant, for the purpose of this experiment, was topical SDS. The inflammatory response was clinically assessed using an established global evaluation scale in which four clinically observable signs of inflammation were assigned an integer score between 0 (“none”) and 3 (“severe”). The four scored endpoints, used to calculate the global score, were: (1) erythema, (2) oedema, (3) skin dryness/desquamation and (4) vesicles. The extract was tested using two different experimental protocols. In the first protocol, SDS was topically applied to the skin in conjunction with the watercress extract. In the second protocol, SDS alone was added to the skin at time t = 24 h, followed by the topical application (24 h later) of the extract. We demonstrated that the watercress extract was able to block the inflammatory response when added either in conjunction with SDS, or when added after induction of the inflammatory response.

The nature of the bioactive compounds that explain the anti-inflammatory activity of the watercress extract, as induced by SDS-mediated skin damage, remains to be elucidated. However, two key classes of molecules, known to be present in watercress, could be linked to a skin anti-inflammatory effect: isothiocyanates and polyphenols, including flavonoids. Both isothiocyanates and flavonoids exhibit anti-inflammatory activities, although the modes of action of these two classes of compounds may be different. Some isothiocyanates, acting as electrophiles, have the capacity to act as potent Nrf2 inducers, thereby exerting anti-inflammatory effects, in skin cells (Gęgotek and Skrzydlewska, 2015). Activation of the transcription factor Nrf2 is involved the upregulation of cytoprotective genes such as NAD(P)H:quinone oxidoreductase 1 (NQO1; Ernst et al, 2011).

The isothiocyanate, sulforaphane, was putatively identified in the present extract, and has been reported by other investigators to be present in watercress (Revelou et al, 2022), albeit at lower concentrations than in some other cruciferous vegetables. Topical application of a sulforaphane-rich broccoli extract was shown to upregulate NQO1 expression and to exert anti-inflammatory activity in various models of skin inflammation, including protection against UV-induced erythema in mouse and human skin (Talalay et al, 2007). Sulforaphane, and other known Nrf2 activating substances, have anti-inflammatory effects in animal models of eczema and psoriasis (reviewed by Ogawa and Ishitsuma, 2022). All four of the isothiocyanates detected in the present extract have been reported to upregulate NQO1 in cultured cells (Rose et al 2000; Ernst et al, 2011). Lim et al (2017) tested the effects of 7-methylsulfinylheptyl isothiocyanate in a cultured macrophage cell line (RAW 264.7) challenged with the pro-inflammatory stimulus, lipopolysaccharide (LPS). The latter isothiocyanate was able to block several inflammatory pathways, including NF-κB activation. Rose et al (2005a) observed similar anti-inflammatory effects of PEITC and 8-methylsulphinyloctyl isothiocyanate in the same cultured cell type. These studies suggest that the mixture of isothiocyanates present in the watercress extract may be involved in its anti-inflammatory activity. However, watercress also contains a wide array of polyphenols which have also been suggested to explain the topical anti-inflammatory activity of watercress extracts (Sadeghi et al, 2013; Camponogara et al, 2019).

A limitation of the present study is that the chemical characterisation of the extract included a limited array of identified compounds. Further chemical analysis will provide a better understanding of the observed bioactivities of the extract. The present report was limited to the putative identification of constituent compounds by LC-MS. The presented compound identities require confirmation by LC-MS/MS, as well the use of authentic standards to allow quantification of the concentrations of the identified compounds. In the present study, a topical anti-inflammatory activity of the extract was observed at a concentration of 10% watercress extract, in the context of SDS-induced human skin inflammation. Concentration-response studies will be essential in the future exploration of the potential beneficial effects of the extract on skin irritation. Camponogara et al (2019) tested the topical application of a crude hydroalcoholic extract of watercress, at a broadly similar concentration (up to 3%), in croton-oil-induced mouse ear oedema. The 3% extract concentration resulted in significant reductions in epidermal thickness and other indices of inflammation.

We plan to test the present extract in other models of skin inflammation. Clearly, our results from the single model of SDS-induced skin inflammation cannot be assumed to translate to anti-inflammatory effects when skin is subjected to other environmental and immune-related inducers of skin inflammation. Nevertheless, the reported possibility of producing a non-toxic aqueous extract of watercress – suitable for application to human skin whilst possessing ant-inflammatory constituents/bioactivity – warrants further investigation.

## 5. Acknowledgements

K.S. and P.G.W. are co-founders of Watercress Research Ltd and are Inventors in relation to patent filings (WO2022129618A1) comprising the preparation and uses of watercress extracts (owned by the University of Exeter, UK). We thank Tom Amery, of The Watercress Company, Dorchester, UK, for organising the supply of fresh watercress.

## Supplementary data

**Supplementary Table 1.**
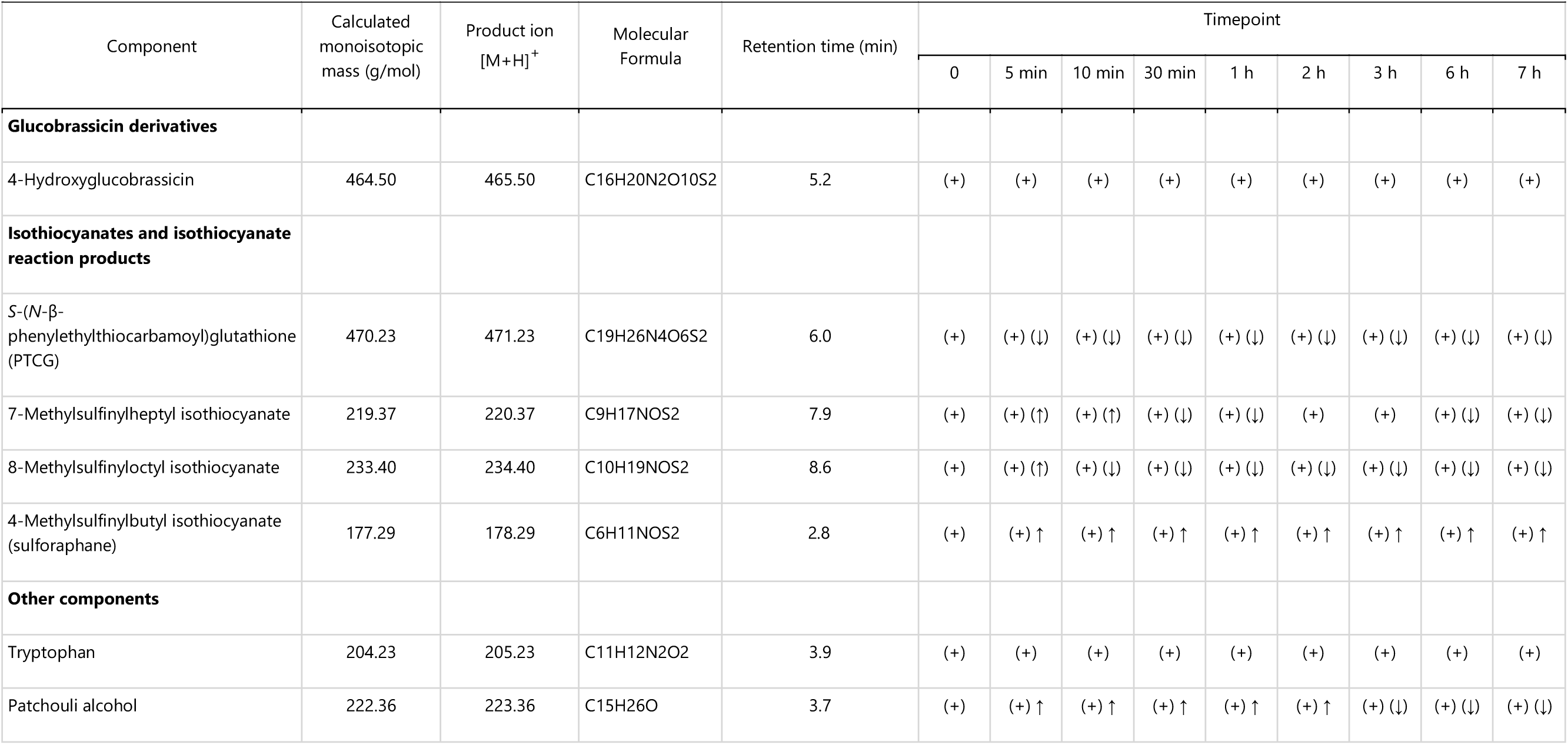
Analysis of watercress aqueous extract – UHPLC-QTOF-MS data generated in positive mode. (+) Tentative assignment, based on chromatographic elution time and mass spectral analysis, (↑) Concentration increasing with time, (↓) Concentration decreasing with time.

| Component | Calculated monoisotopic mass (g/mol) | Product ion [M+H]<sup>+</sup> | Molecular Formula | Retention time (min) | Timepoint |  |  |  |  |  |  |  |  |
| --- | --- | --- | --- | --- | --- | --- | --- | --- | --- | --- | --- | --- | --- |
|  |  |  |  |  | 0 | 5 min | 10 min | 30 min | 1 h | 2 h | 3 h | 6 h | 7 h |
| **Glucobrassicin derivatives** |  |  |  |  |  |  |  |  |  |  |  |  |  |
| 4-Hydroxyglucobrassicin | 464.50 | 465.50 | C16H20N2O10S2 | 5.2 | (+) | (+) | (+) | (+) | (+) | (+) | (+) | (+) | (+) |
| **Isothiocyanates and isothiocyanate reaction products** |  |  |  |  |  |  |  |  |  |  |  |  |  |
| S-(N-β-phenylethylthiocarbamoyl)glutathione (PTCG) | 470.23 | 471.23 | C19H26N4O6S2 | 6.0 | (+) | (+) (↓) | (+) (↓) | (+) (↓) | (+) (↓) | (+) (↓) | (+) (↓) | (+) (↓) | (+) (↓) |
| 7-Methylsulfinylheptyl isothiocyanate | 219.37 | 220.37 | C9H17NOS2 | 7.9 | (+) | (+) (↑) | (+) (↑) | (+) (↓) | (+) (↓) | (+) | (+) | (+) (↓) | (+) (↓) |
| 8-Methylsulfinyloctyl isothiocyanate | 233.40 | 234.40 | C10H19NOS2 | 8.6 | (+) | (+) (↑) | (+) (↓) | (+) (↓) | (+) (↓) | (+) (↓) | (+) (↓) | (+) (↓) | (+) (↓) |
| 4-Methylsulfinylbutyl isothiocyanate (sulforaphane) | 177.29 | 178.29 | C6H11NOS2 | 2.8 | (+) | (+) ↑ | (+) ↑ | (+) ↑ | (+) ↑ | (+) ↑ | (+) ↑ | (+) ↑ | (+) ↑ |
| **Other components** |  |  |  |  |  |  |  |  |  |  |  |  |  |
| Tryptophan | 204.23 | 205.23 | C11H12N2O2 | 3.9 | (+) | (+) | (+) | (+) | (+) | (+) | (+) | (+) | (+) |
| Patchouli alcohol | 222.36 | 223.36 | C15H26O | 3.7 | (+) | (+) ↑ | (+) ↑ | (+) ↑ | (+) ↑ | (+) ↑ | (+) (↓) | (+) (↓) | (+) (↓) |

**Supplementary Table 2.**
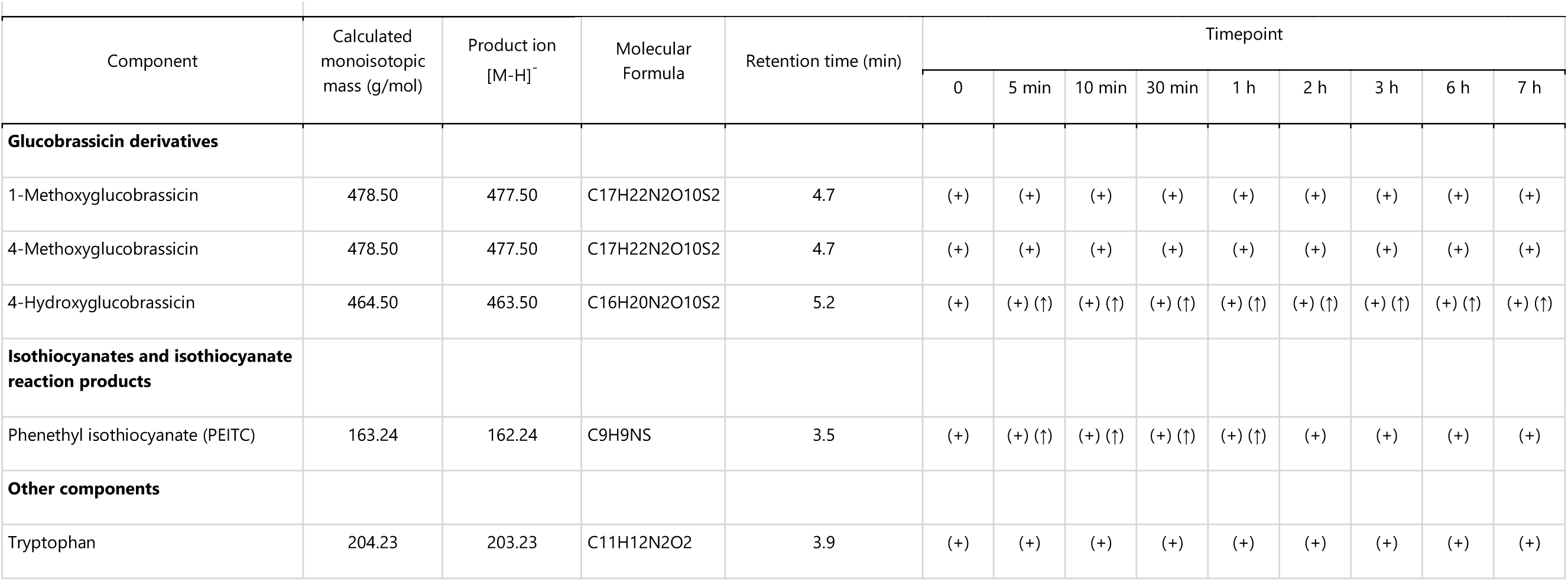
Analysis of watercress aqueous extract – UHPLC-QTOF-MS data generated in negative mode. (+) Tentative assignment, based on chromatographic elution time and mass spectral analysis, (↑) Concentration increasing with time, (↓) Concentration decreasing with time.

| Component | Calculated monoisotopic mass (g/mol) | Product ion [M-H]<sup>-</sup> | Molecular Formula | Retention time (min) | Timepoint |  |  |  |  |  |  |  |  |
| --- | --- | --- | --- | --- | --- | --- | --- | --- | --- | --- | --- | --- | --- |
|  |  |  |  |  | 0 | 5 min | 10 min | 30 min | 1 h | 2 h | 3 h | 6 h | 7 h |
| **Glucobrassicin derivatives** |  |  |  |  |  |  |  |  |  |  |  |  |  |
| 1-Methoxyglucobrassicin | 478.50 | 477.50 | C17H22N2O10S2 | 4.7 | (+) | (+) | (+) | (+) | (+) | (+) | (+) | (+) | (+) |
| 4-Methoxyglucobrassicin | 478.50 | 477.50 | C17H22N2O10S2 | 4.7 | (+) | (+) | (+) | (+) | (+) | (+) | (+) | (+) | (+) |
| 4-Hydroxyglucobrassicin | 464.50 | 463.50 | C16H20N2O10S2 | 5.2 | (+) | (+) (↑) | (+) (↑) | (+) (↑) | (+) (↑) | (+) (↑) | (+) (↑) | (+) (↑) | (+) (↑) |
| **Isothiocyanates and isothiocyanate reaction products** |  |  |  |  |  |  |  |  |  |  |  |  |  |
| Phenethyl isothiocyanate (PEITC) | 163.24 | 162.24 | C9H9NS | 3.5 | (+) | (+) (↑) | (+) (↑) | (+) (↑) | (+) (↑) | (+) | (+) | (+) | (+) |
| **Other components** |  |  |  |  |  |  |  |  |  |  |  |  |  |
| Tryptophan | 204.23 | 203.23 | C11H12N2O2 | 3.9 | (+) | (+) | (+) | (+) | (+) | (+) | (+) | (+) | (+) |

